# The effects of cell-cell orientation in modeling the hallmarks of lung cancer in vitro

**DOI:** 10.1101/2023.12.30.573475

**Authors:** Andres S. Espinoza, Rachael N. McVicar, Darren Finlay, Rabi Murad, Kristiina Vuori, Bethany A. Grimmig, Anne Bush, Emily Smith, Thomas Mandel-Clausen, Heather M. McGee, Evan Y. Snyder, Sandra L. Leibel

## Abstract

To better understand and develop treatments for lung cancer, it is important to have reliable and physiologically relevant culture models. Traditional methods of growing lung cancer cells in two-dimensional (2D) monolayers have limitations in mimicking the complex architecture and microenvironment of lung tumors *in vivo*, limiting their value as reliably informative disease models. In this study, we introduce a new cell culture platform called “tumoroids,” which involves growing HCC827 lung cancer cells in three-dimensional (3D) configurations. By comparing transcriptional profiles of HCC827 cells grown as tumoroids versus the same cells grown in 2D monolayers, we investigate how cell-cell orientation and signaling impact the cancer-driving properties of lung cancer. We examine key features associated with cancer progression, such as epithelial mesenchymal transition, replicative ability, and induction of angiogenesis. Additionally, we assess the functional characteristics of the 3D tumoroid culture system by subjecting the cells to irradiation. Through comparing the gene expression profiles of the 3D tumoroid and 2D cultures with those of primary human lung adenocarcinoma, we find that the 3D tumoroid cultures more closely resemble the characteristics of primary human lung cancer. This suggests that the 3D tumoroid culture platform can serve as a valuable *in vitro* model for studying lung cancer, offering greater clinical relevance compared to traditional 2D cultures. Overall, this study highlights the importance of using advanced culture models like 3D tumoroids to improve our understanding of lung cancer and facilitate the development of effective treatments.

## INTRODUCTION

Lung cancer is a widespread and devastating disease that affects both men and women and is the leading cause of cancer-related deaths worldwide (1). Researchers have been working to improve outcomes by implementing early screening practices and developing targeted therapies, such as anti-EGFR antibodies and immune checkpoint inhibitors (2–3).

One of the challenges in human lung cancer research is recreating the complex environment of tumors *in vitro*. Tumors consist of various types of cells and have a heterogeneous nature, making it difficult to accurately predict how they will respond to treatments (4). Traditional research methods have relied on two-dimensional (2D) epithelial cell lines, patient-derived xenografts (PDX), and genetically engineered mouse models (GEMM) to study lung cancer development and test drug responses (4). However, these models have not been effective in predicting clinical outcomes in patients. To overcome these limitations, investigators have turned to three-dimensional (3D) spheroid/organoid models, which aim to mimic the natural 3D structure of cells found in tumors.

Spheroids/organoids preserve the characteristics of multiple cell types present in the tumor (5).

Most pre-clinical studies investigating lung cancer have traditionally relied on 2D monolayer cultures, which are simplified and do not fully replicate the complex microenvironment of lung tissue (6). While 2D models are advantageous due to their ease of replication and lower cost compared to other culture methods, they do not accurately represent the interactions between tumor cells and their surrounding tissues (5).

*In vivo* models, such as patient-derived xenografts (PDX) and genetically engineered mouse models (GEMM), provide a more realistic assessment of tumor biology but have limitations. These models are expensive, time-consuming to establish, and still do not fully capture the complex interactions between tumors and the surrounding stroma as seen in humans (7–8).

In recent years, 3D models have gained popularity in cancer research (9). These models offer a more authentic representation of the in vivo environment compared to traditional 2D cultures. 3D models have several advantages, including more realistic cell shape and orientation, increased surface area exposed to the culture medium, better cell proliferation, improved drug sensitivity, and more accurate response to growth factors and other stimuli (10–11).

We wanted to investigate if the way cancer cells are grown in different configurations, either as 2D monolayers or 3D “tumoroids,” can alter cancer-related signaling. In lung cancer, there are eight primary hallmarks that drive cancer progression, including oncogene activation, loss of tumor suppressor genes, unlimited ability to replicate, induction of angiogenesis, evasion of apoptosis, metastasis, immune evasion, and cell metabolic reprogramming (12). These hallmarks serve as indicators of aberrant signaling within cells that display distinguishing features of cancer (13). In this study, we created 3D tumoroids of lung cancer cells derived from cells that are traditionally grown as 2D monolayers. We examined how the different growth configurations and cell-cell orientations influenced cancer-related properties, such as gene expression and response to irradiation.

Understanding these fundamental aspects of cancer biology is crucial for identifying potential therapeutic targets and developing effective treatments.

## METHODS

### 2D Culture of the HCC827 cells

The HCC827 cell line (ATCC) was grown in a T-75 flask (Corning) and cultured in 10 mL of RPMI 1640 medium with GlutaMAX supplemented with 10% fetal bovine serum (FBS), 100 U/mL penicillin, and 100U/mL streptomycin (complete media). Cells were grown in an incubator at 37°C and 5% CO_2_. Cells were cultured for 4-6 days before passaging. Cells were passaged once they reached 80% confluency. For passaging, 5 ml of dissociation enzyme, TrypLE was added to the HCC827 cells in a T-75 flask and allowed to incubate at 37 degrees for 12-15 minutes until the cell layer was dispersed. The dissociation was quenched with 10mL of DMEM-F12 with 2% FBS. Cells were dissociated by washing gently with a 10ml pipette and collected into a 15 ml conical tube. The cells were spun down at 400 g for 5 min and the supernatant was discarded. The cells were resuspended in 5mL of fresh and warm complete media and counted. One million cells were passaged into a fresh uncoated T-75 flask with 10mL of complete media.

### 3D Culture of the HCC827 cells

To make 3D tumoroids, HCC827 monolayer cells were passaged into growth factor reduced (GFR) Matrigel droplet (Catalog #354230). After monolayer dissociation, the cells were pelleted after spinning down at 400 g for 5 min. The supernatant was discarded, and the cells were resuspended in 5mL of complete media and 50,000 cells were aliquoted into 1.5 ml centrifuge tubes. The tubes were spun down the supernatant discarded. 100 µL of GFR matrigel was added to the cells on ice, resuspended carefully to avoid bubbles, and placed into a 12-well plate as a droplet. The plate was placed in the incubator for gelling to occur for 30 min. 1 mL of 1640 medium with Glutamax supplemented with 10% fetal bovine serum (FBS), 100 U/mL penicillin, and 100U/mL streptomycin was added to each well. The 3D tumoroids embedded in Matrigel were maintained in an incubator at 37°C and 5% CO_2_ for 12-16 days or until they were 80% confluent and fed every 2-3 days.

### 3D Culture Passaging Protocol/ Single Cell Dissociation Protocol

To passage confluent tumoroids onto a new Matrigel droplet, fresh Matrigel was cool thawed on ice into a liquid. The cell media was aspirated from the well and 1mL of Dispase (+Rock Inhibitor (Tocris REF 1254)) was added to each well. The cells were incubated at 37°C for 30 minutes and triturated at 15-minute increments. Dissociated 3D tumoroids were transferred into a new 15mL conical and chilled PBS was added to wash down and “melt” the Matrigel. The conical was spun down at 400 x g for 5 minutes and a P-1000 pipette was used to remove the supernatant. If residual “gray Matrigel” was present, the PBS step was repeated. The Matrigel-free cells were then spun down at 400 x g for 5 minutes and the supernatant was discarded. 1-2mL of TrypLE was added and the cells were incubated at 37°C for 15 minutes. The reaction was quenched by adding 2x volume of stop media (DMEM-F-12 with 2% FBS). The cells were centrifuged at 400 x g for 5 minutes and resuspended in 1mL of complete media and counted. After calculating the number of cells required for a seeding of 50,000 cells per Matrigel drop, the cells were transferred into 1.5mL centrifuge tubes and spun down at 400 x g for 5 minutes. Supernatant was removed and the cells were carefully resuspended in 100uL of Matrigel and transferred into a 12-well plate as a droplet. The plate was placed in the incubator for gelling to occur for 30 min. After the droplets have gelled, 1 mL of complete media was added to each droplet.

### Western Blot

3D and 2D HCC827 cells were lysed with RIPA lysis buffer supplemented with protease inhibitors and phosphatase inhibitors (HALT kit, Thermo Scientific). Lysates were centrifuged at 16,000 x g at 4 °C for 25 minutes and the supernatant was collected. A BCA (Thermo Scientific, REF23225) was conducted to determine protein concentration. 10-20ug of protein was mixed with NuPage sample buffer (Life Technologies, REF NP0007) supplemented with dithiothreitol 100uM then were heated at 95°F for 5 minutes. They were then run through SDS-PAGE with MES running buffer (Life Technologies, REF NP0002) for 45 minutes and 80 mAmps and then transferred to a 45uM PVDF membrane (Thermo Scientific REF 88518). The transfer was run using the NuPage transfer buffer for 90 minutes at 20 volts. The membrane was blocked in TBST with 5% non-fat milk diluted in TBS with 0.1% tween-20 (TBST) for one hour at room temperature. Primaries were diluted in 1% non-fat milk in TBS-T incubated overnight at 4°C. Secondary antibodies (LICOR goat anti rabbit 800CW, 925-32213, and goat anti mouse 680, 925-68072, both diluted at 1:10;000) were added the following day for one hour at room temperature. The western blot analysis was conducted using the Odyssey near infra-red scanner to quantify pixel intensity of the designated bands, data was later compiled using Excel and statistics run using Prism 8 software.

### Immunohistochemistry

2D monolayer HCC827 cells were dissociated as previously described in the 2D cell culture protocol, then 200,000 cells plated per well of an 8-well glass chamber slide. Once the HCC827 monolayer reached 80% confluency, the cells were fixed with 4% Paraformaldehyde (PFA) at room temp for 15 minutes, then washed with PBS three times. The samples were blocked for one hour at room temperature with staining buffer composed of 5% BSA, 5% Donkey serum, 0.2%Tri-X 100 in PBS -/-. Then, the 2D monolayer was stained with primary antibodies at company recommended concentrations, diluted in staining buffer, at 4°C overnight. 100ul staining volume per well was used. The following day, washes were performed 3x with PBS -/-, then secondary antibodies were added for one hour at room temperature. Samples were covered to prevent photobleaching from light. 3x washes were performed after secondary, and nuclei stained with Hoechst (Invitrogen #H3570) was added at 1:5000 for 15 minutes. Samples were washed 3x with PBS -/-, then mounted with cover glass and FlouroSave reagent (Millipore# 345789). The figures were processed using ImageJ software.

Once the HCC827 3D tumoroids reached confluency (usually 1-2 weeks post-Matrigel embedding), the tumoroid-containing Matrigel sample was fixed with 4% Paraformaldehyde for one hour at room temperature and washed with PBS 3x. Fixed tumoroids in Matrigel were processed by the Sanford Burnham Histology core, where samples were embedded in paraffin wax, sliced, and mounted onto glass slides. On the day of staining, antigen retrieval was performed on paraffin sectioned slides. Staining was performed as indicated for the 2D samples.

Samples were imaged using the Zeiss LSM 780 Confocal Microscope and the Olympus IX71/IX81 IX2 series Inverted Phase Contrast/Fluorescence Microscope

### Quantification of immunofluorescence

Images for each nuclear marker were quantified using ImageJ. Briefly, images were converted to 8-bit images and the threshold was adjusted to correspond with the nuclear stain, which allows for measurement of total area. The total area was analyzed by the Analyze Particles” function of ImageJ. Percentage of positive cells were calculated by dividing the total area of positive cells over the total area of DAPI. For extracellular matrix quantification, fluorescence intensity was quantified using Leica Application Suite X.

### RNA Isolation

Total RNA from the 2D and 3D HCC827 cell lines was purified using the *Direct-zol RNA MicroPrep kit (Genesee Scientific: 11-330MB)* using the manufacturer’s recommendations, including DNase treatment. RNA concentration and RNA integrity number (RIN) were determined using an Agilent microfluidic RNA 6000 Nano Chip kit (Agilent Technologies, Santa Clara, CA) on the 2100 Bioanalyzer (Agilent Technologies, Santa Clara, CA). Those samples with RIN greater than 9 were used for RNAseq. PolyA RNA was isolated using the NEBNext® Poly(A) mRNA Magnetic Isolation Module and barcoded libraries were made using the NEBNext® Ultra II™ Directional RNA Library Prep Kit for Illumina® (NEB, Ipswich MA). Libraries were pooled and single end sequenced (1X75) on the Illumina NextSeq 500 using the High output V2 kit (Illumina Inc., San Diego CA). Read data was processed in BaseSpace (basespace.illumina.com). Reads were aligned to Homo sapiens genome (hg19) using STAR aligner (https://code.google.com/p/rna-star/) with default settings.

### Bulk RNA Seq Analysis

Illumina sequencing reads were preprocessed to remove Truseq adapter, polyA, and polyT sequences using cutadapt v2.3 (Martin 2011). Trimmed reads were aligned to human genome version 38 (hg38) using STAR aligner v2.7.0d_0221 (Dobin, Davis et al. 2013) with parameters according to ENCODE long RNA-seq pipeline (https://github.com/ENCODE-DCC/long-rna-seq-pipeline). Gene expression levels were quantified using RSEM v1.3.1 (Li and Dewey 2011) and Ensembl gene annotations version 84. RNA-seq sequence, alignment, and quantification quality was assessed using FastQC v0.11.5 (https://www.bioinformatics.babraham.ac.uk/projects/fastqc/) and MultiQC v1.8 (Ewels, Magnusson et al. 2016). Biological replicate concordance was assessed using principal component analysis (PCA) and pairwise pearson correlation analysis. Lowly expressed genes were filtered out by applying the following criterion: estimated counts (from RSEM) ≥ number of samples * 5. Filtered estimated read counts from RSEM were compared using the R Bioconductor package DESeq2 v1.22.2 based on generalized linear model and negative binomial distribution (Love, Huber et al. 2014). Genes with Benjamini-Hochberg corrected p-value < 0.05 and fold change ≥ 2.0 or ≤ -2.0 were selected as differentially expressed genes. Differentially expressed genes were then analyzed using Ingenuity Pathway Analysis (Qiagen, Redwood City, USA)

### Extraction of intact 3D Cells from Matrigel

1mL of Cell Recovery Solution (Catalog #354253) was added to the matrigel droplets. The tip of a P1000 was trimmed to prevent lysis of tumoroids. Using the cut down P1000, the tumoroids were resuspended in the solution, transferred to a 15mL conical and incubated in 4 degrees for 1 hour.

Cells were gently agitated by flicking the tube every 15 minutes. Cells were then resuspended in 3mL of cold PBS. Samples were centrifuged for 1 minute at 200 x g. Gray-like Matrigel and supernatant was removed using a pipette. 1 mL of complete culture media was added, and a sample was taken for counting.

### Irradiation

2D monolayer cells were dissociated as indicated in previous dissociation protocol and approximately 20,000 cells were plated in 2 96-well plates in 100uL of culture medium (one control plate and one irradiation plate). The 3D tumoroids were extracted from Matrigel and approximately 20,000 cells were plated in 2 ULA 96-well plates (one control plate and one irradiation plate). 100uL of CellTiter-Glo Reagent (Promega Corp., Catalog # G7572) was used to lyse the cells and measure cell death at Timepoint 0 for the non-irradiated 2D and 3D HCC827 plates. 2D and 3D irradiation plates were then exposed to a dose of 20 Gy of irradiation using the RadSource RS 2000. All 4 plates were then incubated at 37°C and cell death was subsequently assessed at the 0, 24, and 72 hour timepoints of both the non-irradiated and the irradiated plates.

### Statistical Analysis

ICC analysis was conducted using the ImageJ software. Western blot bands were recorded in ImageStudioLite and analysis was conducted using Excel and Prism 8 software.

Statistical analysis was performed with Prism 8 (GraphPad Software). We compared all control and irradiation time course data by two-way analysis of variance (ANOVA) followed by a Šidák multiple comparisons test.

## RESULTS

### The lung adenocarcinoma cell line (HCC827) can be grown in 2D and 3D cell culture conditions

Non-small cell lung cancer (NSCLC) is a type of lung cancer that includes two main types: squamous cell carcinoma and adenocarcinoma. These cancers arise from epithelial cells found in the bronchi and extend to the alveoli of the lungs. In adenocarcinoma, a subset of tumors (10-15%) have a specific mutation in the Epidermal Growth Factor Receptor (EGFR) gene (14). When activated, EGFR triggers intracellular signaling pathways that regulate cell growth and survival (15). Treatment for EGFR mutant lung cancers often involves targeted therapies known as tyrosine kinase inhibitors, depending on the specific location of the mutation. The cells of origin for lung adenocarcinoma are believed to be club cells and alveolar type II (ATII) cells (16). ATII cells play important roles in the lungs by secreting surfactant, which helps reduce surface tension and prevent lung collapse. They also self-renew and give rise to alveolar type I (ATI) cells.

In this study, the NSCLC cell line HCC827 was used to investigate the differences between growing the same cancer cells in 2D and 3D culture systems. The HCC827 cell line carries specific mutations in the EGFR gene, including a deletion in Exon 19 (E714 – A750 deletion) and a missense mutation of T790M in Exon 20 (Figure 1A, Supplementary Figure 1). To establish the 2D culture system, the cells were plated in a flask and allowed to grow for 5-7 days. When they reached 80% confluence, the cells were passaged to reduce cell death and stress (Figure 1B). For the 3D culture system, the cells from the monolayer were dissociated, and mixed with chilled liquid Matrigel. This mixture was then placed as droplets in a culture plate and allowed to gel at 37°C. Over time, the cells self-organized into round structures called “tumoroids” within the Matrigel droplets (Figure 1B). The tumoroids proliferated rapidly and protruded single cells out of the Matrigel within 5 days. These cells formed a monolayer outside of the Matrigel, as confirmed by staining with DAPI (Figure 1C). These cells were discarded and not utilized in the analysis of 3D tumoroid samples. The proliferation of cells was assessed by examining the expression of Ki-67, a marker of cellular proliferation (Figure 1D).

**Figure 1:**
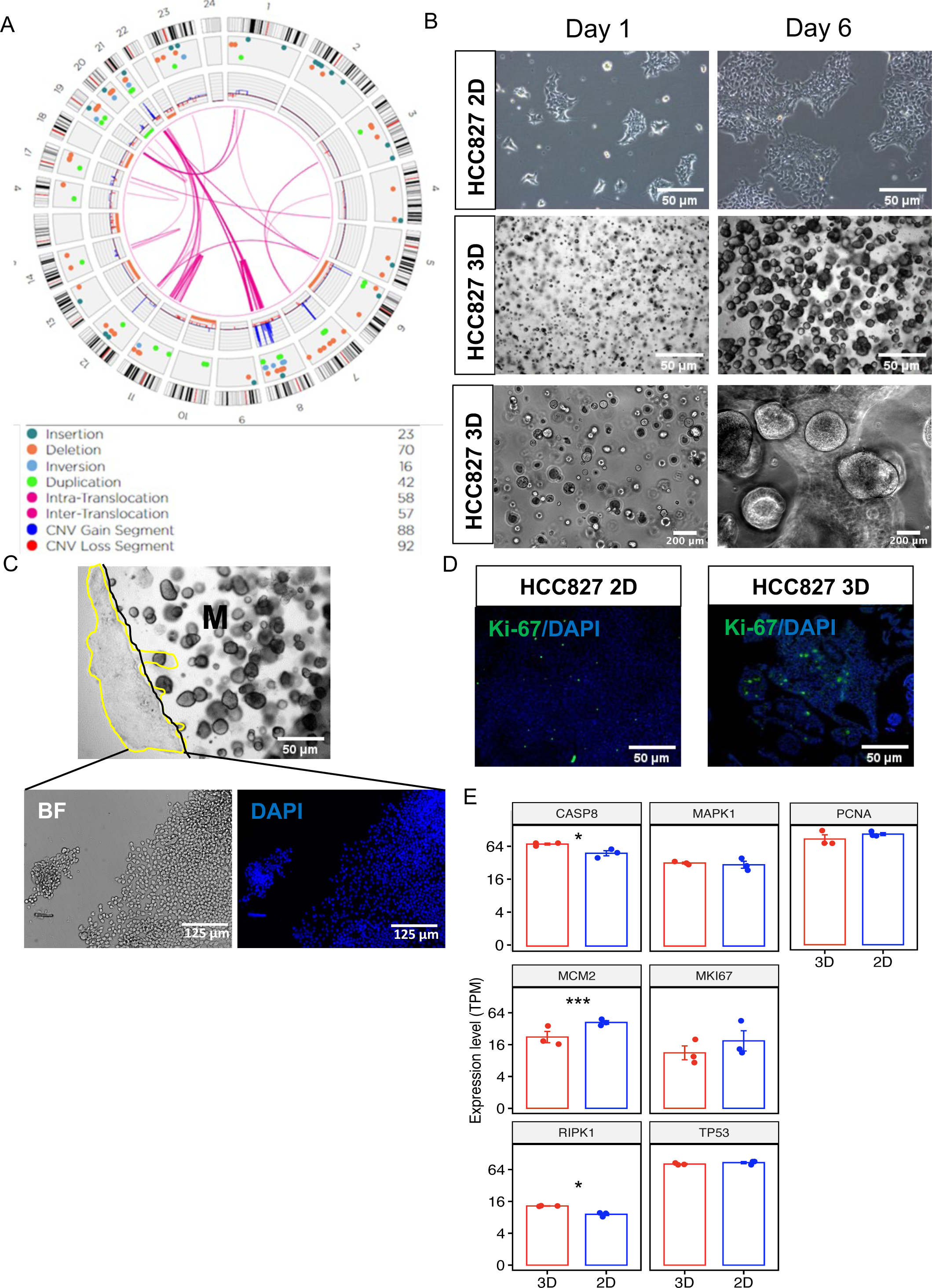
HCC827 adenocarcinoma NSCLC line in 3D vs. 2D cell culture conditions. (a) Schematic of the chromosome map of the adenocarcinoma cell line HCC827. The EGFR mutation is seen in chromosome 7, indicated by the blue spike. **(b)** Brightfield images of live cell cultures in 2D and 3D orientations at days 1 and 6. **(c)** Brightfield images of live cell 3D tumoroid cultures with outgrowth of single cells from the Matrigel (outlined in yellow). Edge of Matrigel (M) outlined in black. **Insets** are a close-up view of these cells in the periphery under brightfield (BF) microscopy and after DAPI nuclear staining. Insets are 2.5 x original images. **(d)** Immunostaining of HCC827 cells in 2D vs. 3D for the proliferation marker Ki-67. The data are representative of at least three independent experiments. A DAPI nuclear stain (blue) is used to visualize all cells in the field. **(e)** Transcript per million (TPM) of growth-associated and death-associated genes in both 3D and 2D cell cultures. (* p<0.05; ** p<0.01; *** p<0.001).

Additionally, we compared the expression of genes related to growth and cell death between the 2D and 3D culture systems (Figure 1e).

### Gene expression profiles differ between the same HCC827 lung cancer cells grown in 2D vs. **3D cell culture conditions.**

To investigate the impact of cell orientation on gene expression profiles, we conducted transcriptional profiling of the HCC827 cells in both 2D and 3D culture environments using bulk RNA sequencing. Principal Component Analysis (PCA) revealed a clear separation between the HCC827 cells grown in 2D and 3D along the first principal component (PC1) (Figure 2A). Differential gene expression analysis comparing the 3D and 2D cultures identified 1568 significantly altered genes (450 downregulated, 1118 upregulated) (Figure 2B). The gene with the highest fold change in the 3D cultures was *BPIFA1*. To gain insights into the functional implications of these gene expression changes, we performed gene set enrichment analysis (GSEA). This analysis revealed several pathways that were significantly upregulated in the 3D culture compared to the 2D culture, including Tumor microenvironment, HIF1α Signaling, Cancer Metastasis Signaling, IL-17 Signaling, ILK Signaling, Regulation of EMT by Growth Factors Pathway, and ERK/MAPK Signaling pathways. On the other hand, the Inhibition of Matrix Metalloproteases Pathway was significantly downregulated (Figure 2C). Furthermore, we examined pathways associated with cancer hallmarks and found differences between the 3D and 2D culture conditions. Genes involved in HIF1α signaling, regulation of epithelial-mesenchymal transition (EMT), and the tumor microenvironment pathway (TMP) were upregulated in the 3D tumoroids compared to the 2D monolayers (Figure 2D). This prompted us to conduct further analysis of these three pathways.

**Figure 2:**
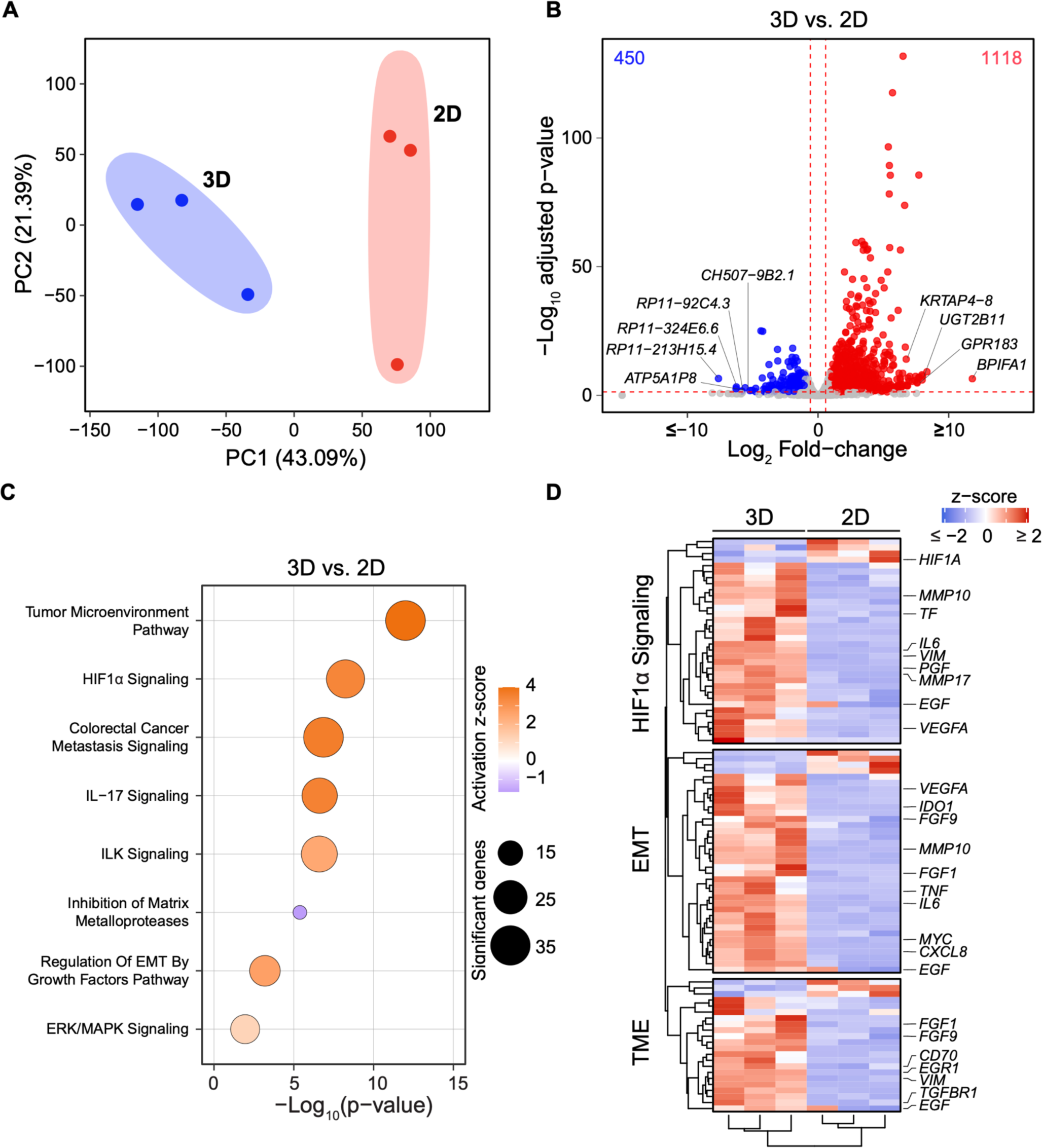
The gene expression profiles differ between HCC827 2D vs 3D culture systems. (a) Principal component analysis (PCA) plot of HCC827 cells in 3D vs. 2D. Each data point indicates an independent replicate (n=3). **(b)** Volcano plot analysis of differential expression of HCC827 cells in 3D vs. 2D systems. The red lines indicate a p-adjusted value <0.05. **(c)** Gene Enrichment Pathways activated and inhibited between HCC827 cells cultured in 3D vs. 2D. The size of the circles represents significant genes, and the color gradient represents the activation z-score. **(d)** Heatmap analysis showing average expression (represented as z-score) of genes important in the HIF1a signaling pathway, Regulation of EMT pathway, and Tumor Microenvironment Pathway in HCC827 cells cultured in 3D vs. 2D. The majority are regulated in the 3D culture system (see text for details).

### Tumor microenvironment pathway genes and proteins are upregulated in the HCC827 3D **culture system.**

The tumor microenvironment (TME) plays a crucial role in cancer progression, including angiogenesis, invasion, and metastasis. It consists of various cell types, including epithelial cancer cells, stromal cells, immune cells, and components of the extracellular matrix (ECM) (17). Signals from tumor cells recruit stromal cells to promote tumorigenicity (18). Stromal cell reprogramming is modulated by cytokines, chemokines, and growth factors secreted by cancer cells (19). Enzymes are also released that degrade and reshape the ECM to facilitate tumor invasion (Figure 3A). Cancer cells recruit non-malignant support cells and induce gene expression changes that promote tumor growth, resistance to apoptosis, angiogenesis, and metastasis. In our study using the HCC827 cell line, which lacks endothelial or immune cells, we focused on understanding gene expression changes and cell signaling within malignant epithelial cells based solely on their configuration in the 3D system.

**Figure 3:**
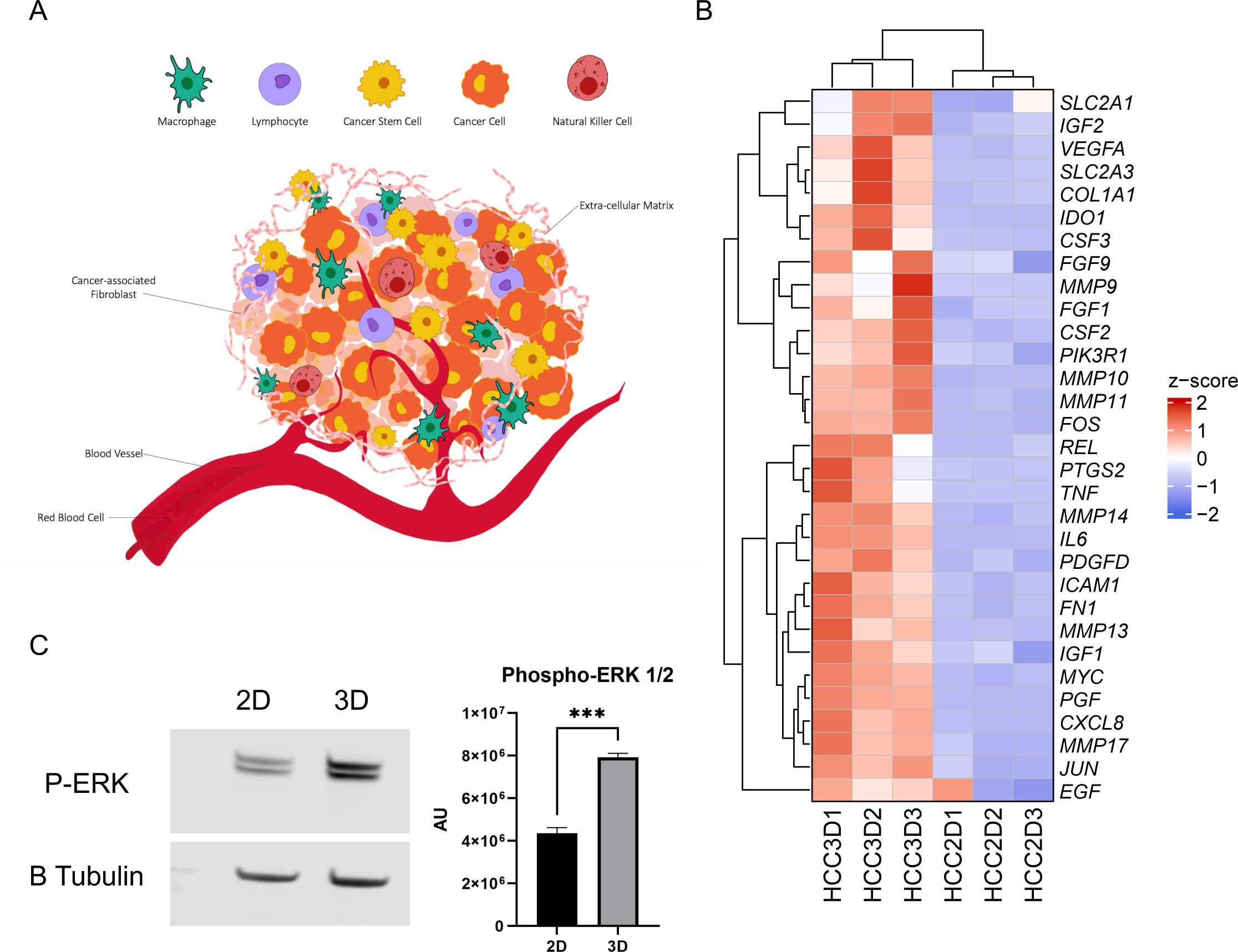
Expression of tumor microenvironment (TME) genes and proteins differ between HCC827 cells cultured in 3D vs. 2D. (a) Schematic of the tumor microenvironment. **(b)** Heatmap analysis showing average expression (represented as z-score) of consistent genes important in the Tumor Microenvironment Pathway in HCC827 cells grown in 3D vs. 2D. Red indicates up-regulated genes and blue indicates down-regulated genes. **(c)** Western blot of phosphorylated extracellular regulated kinase (p-ERK) expression in HCC827 cells in 3D vs. 2D cultures (left panel). Quantification by densitometry (arbitrary units of these cultures shows the that the same cells in 3D evince increased expression of p-ERK compared to the 2D cell cultures (right panel). N=3 biological replicates. **\*\*\*** p<0.001. Error bars in SEM.

Among the pathways examined, the tumor microenvironment pathway exhibited the highest level of activation in the 3D culture system. Genes involved in tumor migration, invasion, metastasis (*MMP9, MMP10, MMP11, MMP13, MMP14, MMP17, ICAM1*), pro-angiogenic factors (*VEGFA, EGF, PDGFD, FGF1, FGF9, IGF1, IGF2*), proliferation genes (*MYC*), and stromal genes (*COL1A1*) were significantly upregulated in the HCC827 3D culture system (p<0.05) (Figure 3B).

To identify potential upstream regulators of these signaling pathways, we evaluated the expression of phosphorylated extracellular signal-regulated kinase 1/2 (ERK1/2), a key molecule in the mitogen-activated protein kinase (MAPK) pathway. Phosphorylated ERK1/2 is an indicator of activated MAPK signaling, which regulates transcription factors involved in proliferation, differentiation, and apoptosis (20). Western blot analysis revealed a significant increase in phosphorylated ERK1/2 in the 3D samples compared to the 2D samples (Figure 3C).

### Hypoxia inducible factor 1 alpha (HIF1**α**) signaling pathway is differentially expressed in the **HCC827 2D and 3D culture systems.**

The increased metabolism and proliferation rates of cancer cells require sufficient access to nutrients for their uncontrolled growth (21). However, tumors often outgrow their blood supply, leading to hypoxia, a state of low oxygen levels. In response to hypoxia, the hypoxia-inducible factor 1 alpha (HIF1α) signaling pathway is activated. HIF1α regulates various downstream effects, including increasing oxygen delivery by promoting erythropoiesis and angiogenesis, reducing oxygen consumption through anaerobic metabolism, and regulating proliferation and apoptosis (22–24).

Additionally, tumors secrete vascular endothelial growth factor (VEGF), a potent angiogenic factor that stimulates the generation of new blood vessels to sustain blood supply to the tumor (Figure 4A).

**Figure 4:**
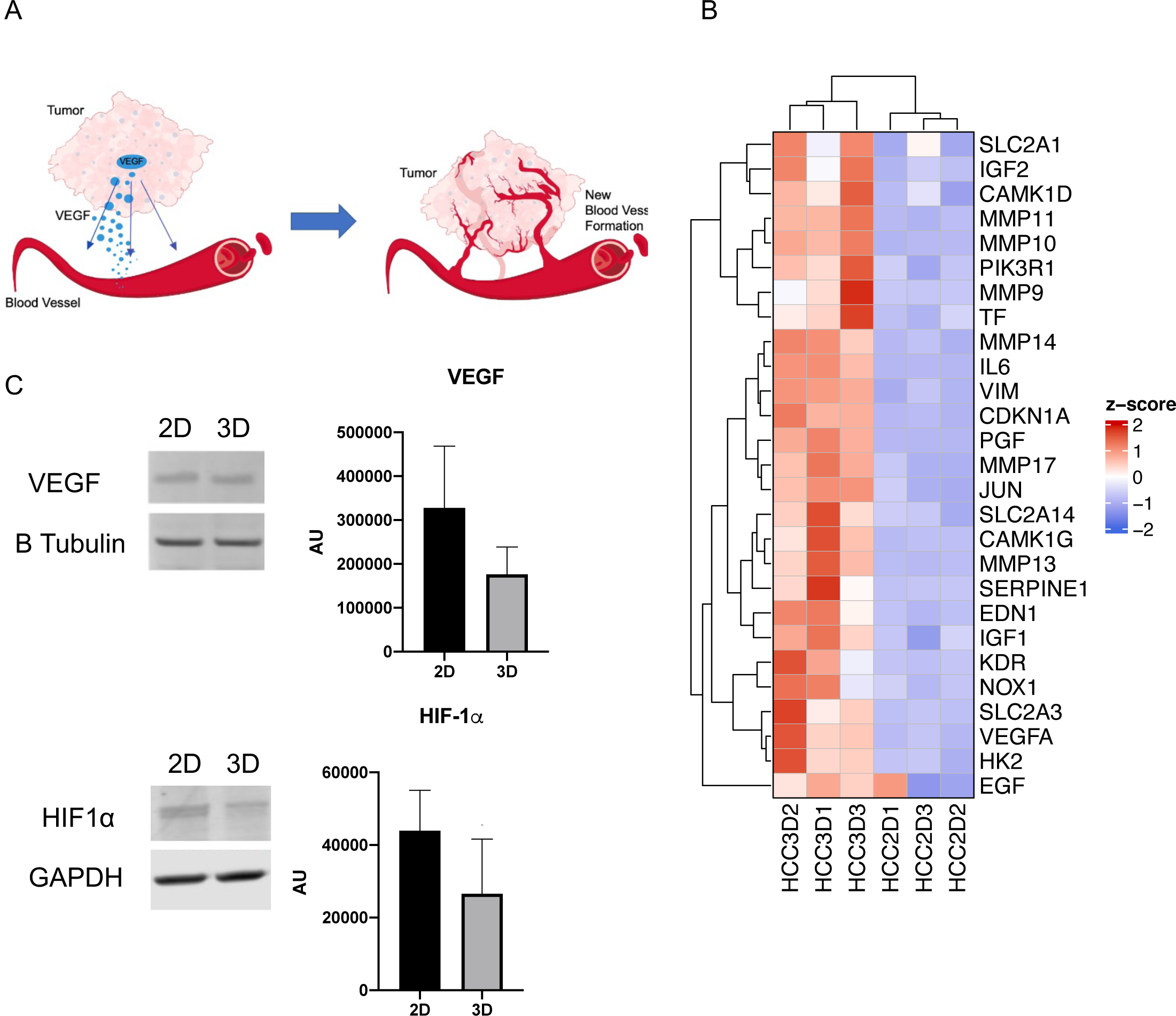
Expression of genes in the tumor angiogenesis signaling pathway genes are differentially expressed in HCC827 cells grown in 3D vs. 2D. (a) Schematic of tumor angiogenesis. **(b)** Heatmap analysis showing average expression (represented as z-score) of consistent genes important in the Angiogenesis Signaling Pathway in HCC827 cells grown in 3D vs. 2D. Red indicates up-regulated genes and blue indicates down-regulated genes. **(c)** Western blots of VEGF (left panel) and HIF1α (right panel) expression by HCC827 cells in 3D vs. 2D cultures. Quantification by densitometry (arbitrary units [AU]) of the protein expression shows no significant differences between the two cell culture systems, although both proteins showed increased expression. N=3 biological replicates. p>0.05. Errors bars in SEM.

In our study, gene expression analysis revealed differences in the HIF1α signaling pathway between the HCC827 2D and 3D cell culture systems. The 3D cell culture system exhibited upregulation of genes involved in this pathway compared to the 2D system. Heatmap analysis confirmed these findings, showing increased activation of *VEGFA* (angiogenesis), *TF* (iron metabolism), and *CDKN1A* (regulation of proliferation and apoptosis) in the 3D cell cultures (Figure 4B). Interestingly, the upstream regulator HIF1α decreased in the 3D model system compared to the 2D system. To validate this finding, we examined the protein expression of VEGF and HIF1α using western blotting. No significant differences were observed in the protein expression of VEGF and HIF1α between the 3D and 2D samples of HCC827 cell cultures (p>0.05) (Figure 4C). Nevertheless, genes involved in the HIF1α signaling pathway and associated with blood vessel formation were significantly upregulated in the 3D tumoroids, including downstream genes regulated by HIF1α.

### Epithelial Mesenchymal Transition (EMT) genes and proteins are up regulated in the HCC827 **3D culture system.**

Epithelial to mesenchymal transition (EMT) is an important hallmark of cancer. During EMT, epithelial cells lose their characteristic cell-cell contacts and polarity, transitioning into a mesenchymal phenotype that enhances cell motility and invasiveness (Figure 5A).

**Figure 5:**
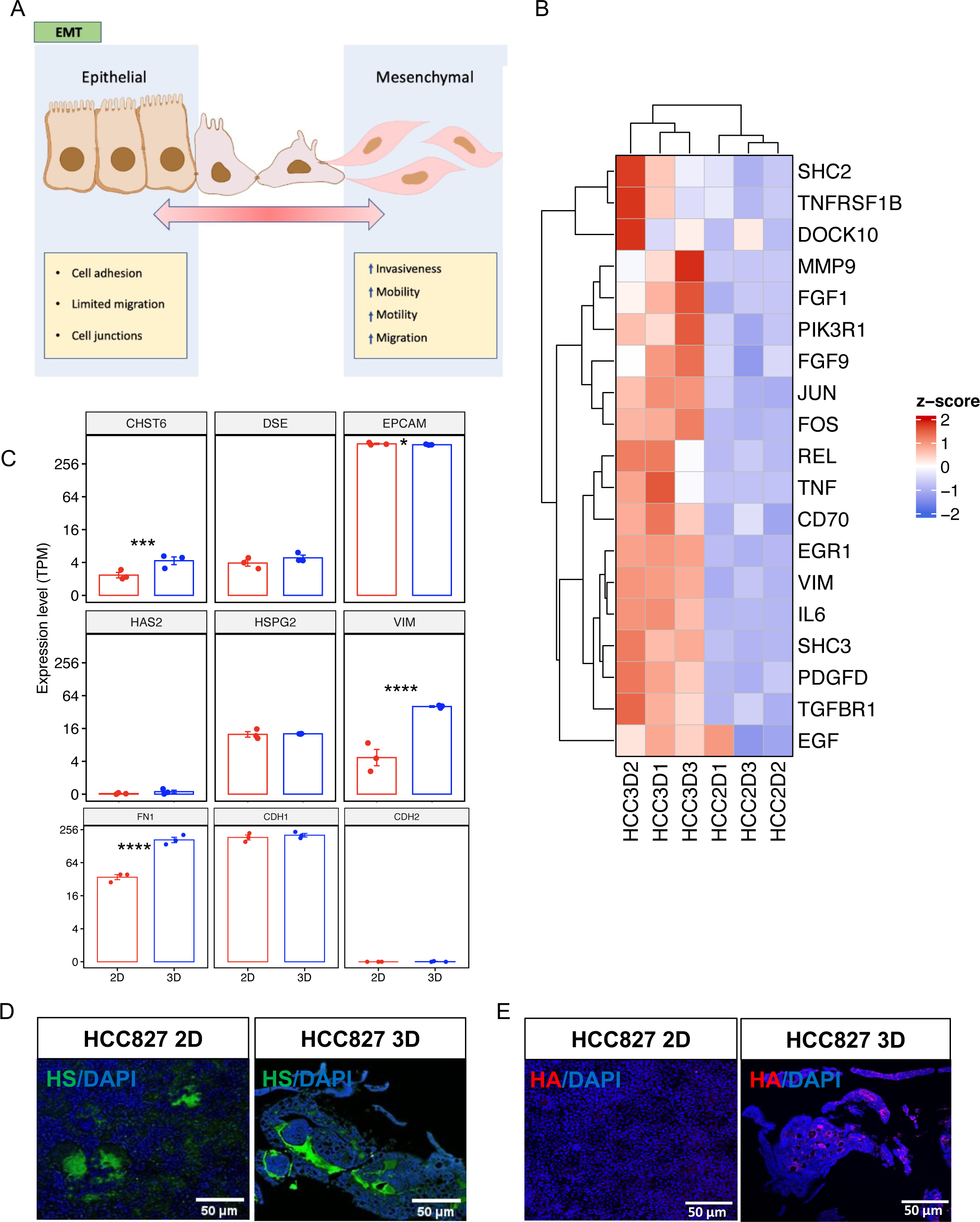
Expression of Epithelial Mesenchymal Transition (EMT) genes and proteins differ in HCC827 cells grown in 3D vs. 2D. (a) Schematic of EMT in cancer cells. **(b)** Heatmap analysis showing average expression (represented as z-score) of consistent genes important in the EMT signaling pathway in HCC827 cells in 3D vs. 2D. Red indicates up-regulated genes and blue indicates down-regulated genes. **(c)** TPM of EMT-associated genes in HCC827 cells in 3D vs. 2D. **(d)** Immunostaining for heparan sulfate (HS) in HCC827 cells in 2D vs. in 3D. **(e)** Immunostaining for hyaluronic acid (HA) in 2D vs. 3D HCC827 cells. The data are representative of at least three independent experiments. A DAPI nuclear stain (blue) is used to visualize all cells in the field. Immunofluorescence was quantified. Quantification of fluorescence showed no differences in HA and HS expression between 3D and 2D HCC827 cells. * p<0.05; *** p<0.001; ****p<0.0001. Errors bars in SEM.

Analysis of gene expression from the bulk RNA sequencing data demonstrated enrichment for EMT in the HCC827 3D cell culture system. Genes associated with mesenchymal cells, such as *VIM* and *MMP9*, were upregulated in the 3D system (Figure 5B). This finding was further supported by TPM analysis, which revealed increased activation of *VIM* and decreased expression of *EPCAM*, an epithelial cell marker, in cells cultivated in the 3D conditions (Figure 5C).

Aberrant glycan expression has been implicated in cancer progression and EMT (25–26). To investigate whether changes in glycan expression were present in the 2D and 3D systems, we examined the expression of heparan sulfate (HS) and hyaluronic acid (HA). These glycans are components of the extracellular matrix (ECM), and their aberrant expression is observed in various types of cancer. HA binds to tumor receptors and induces signaling that promotes cancer cell malignancy (27). High levels of HA have been detected in lung, prostate, ovarian, and breast cancers (28). Similarly, HS can influence tumor phenotype depending on the proteoglycan subset (29). HS has also been shown to bind mitogenic and angiogenesis-promoting growth factors, thereby enhancing carcinogenesis (30).

Gene expression analysis revealed similar expression levels of HS (*HSPG2*) and HA (*HAS2*) in both the 2D and 3D culture systems (Figure 5D). Immunostaining analysis also revealed similar HS and HA protein expression levels (Figure 5E).

### Functional changes in the Surfactant system between the 2D and 3D HCC827 cell culture **systems.**

Pulmonary surfactant, which is secreted by ATII cells in the lung, consists of four proteins: surfactant protein A, B, C, and D (SP-A, -B, -C, and -D, respectively). SP-A and SP-D are hydrophilic proteins involved in regulating innate immune cells in the lung, while SP-B and SP-C are hydrophobic proteins that play a role in reducing surface tension in the lung (31–32). SP-B, in particular, is responsible for facilitating lipid adsorption into the air-liquid interface and stabilizing the multilayer interfacial films during expiration (33–34).

In lung cancer studies, higher protein expression of SP-B has been reported in the blood of genetically engineered lung cancer mouse models and lung cancer patients, suggesting its potential utility as a biomarker for lung cancer (35). Additionally, it has been hypothesized that SP-B could play a role as a suppressor of non-small cell lung cancer (NSCLC) progression (36) (Supplementary figure 2A). Improved outcomes in irradiation therapy have also been observed in an *in vivo* mouse xenograft model after administration of SP-B (36).

To assess the expression of surfactant proteins in the 2D vs. 3D cell culture systems, the expression of all surfactant genes was examined. Among them, only *SFTPA1, SFTPA2*, and *SFTPB* showed significant differences between the model systems (Supplementary figure 2B). The expression levels of their transcription factors, *NKX2-1* and *FOXA2,* were also evaluated upstream, but no difference was found in *NKX2-1*, while *FOXA2* expression was reduced in the 3D model.

Furthermore, the protein expression of surfactant protein B (SP-B) was assessed using western blotting and immunofluorescence. Two isoforms of SP-B were detected in the cells, but only the 42 kDa isoform showed a significant difference between the model systems, elevated in the 3D tumoroids (Supplementary figure 2C).

To confirm the protective role of SP-B in cell injury, scratch assays were performed in the 2D HCC827 culture system in the presence and absence of a whole lung surfactant emulsion containing phospholipids and SP-B (poractant alfa). The scratch assay involved creating a linear injury through the center of a confluent well, and the size of the negative space was compared over 14 hours with and without surfactant. Although the addition of the surfactant trended towards improved healing, the result was not significant (Supplementary Figure 3).

### Response to irradiation differs between the 2D and 3D HCC827 cell culture systems

To assess whether a 3D configuration of lung cancer cells could better predict their sensitivity to a given therapy compared to traditional methods, a proof-of-concept study was conducted using HCC827 cells and irradiation as the therapeutic intervention. Two experimental conditions were employed: 1) HCC827 cells were seeded as monolayers in a 96-well culture plate. 2) 3D tumoroids of HCC827 cells were suspended in the wells of an ultra-low attachment (ULA) 96- well plate. Both sets of cells were then subjected to irradiation at a dosage of 20 Gy using the RadSource RS 2000. Cell viability before and after irradiation was assessed using an ATPase luciferase assay kit, with the luminescence emitted by the wells serving as a measure of cell number (Figure 6A).

**Figure 6:**
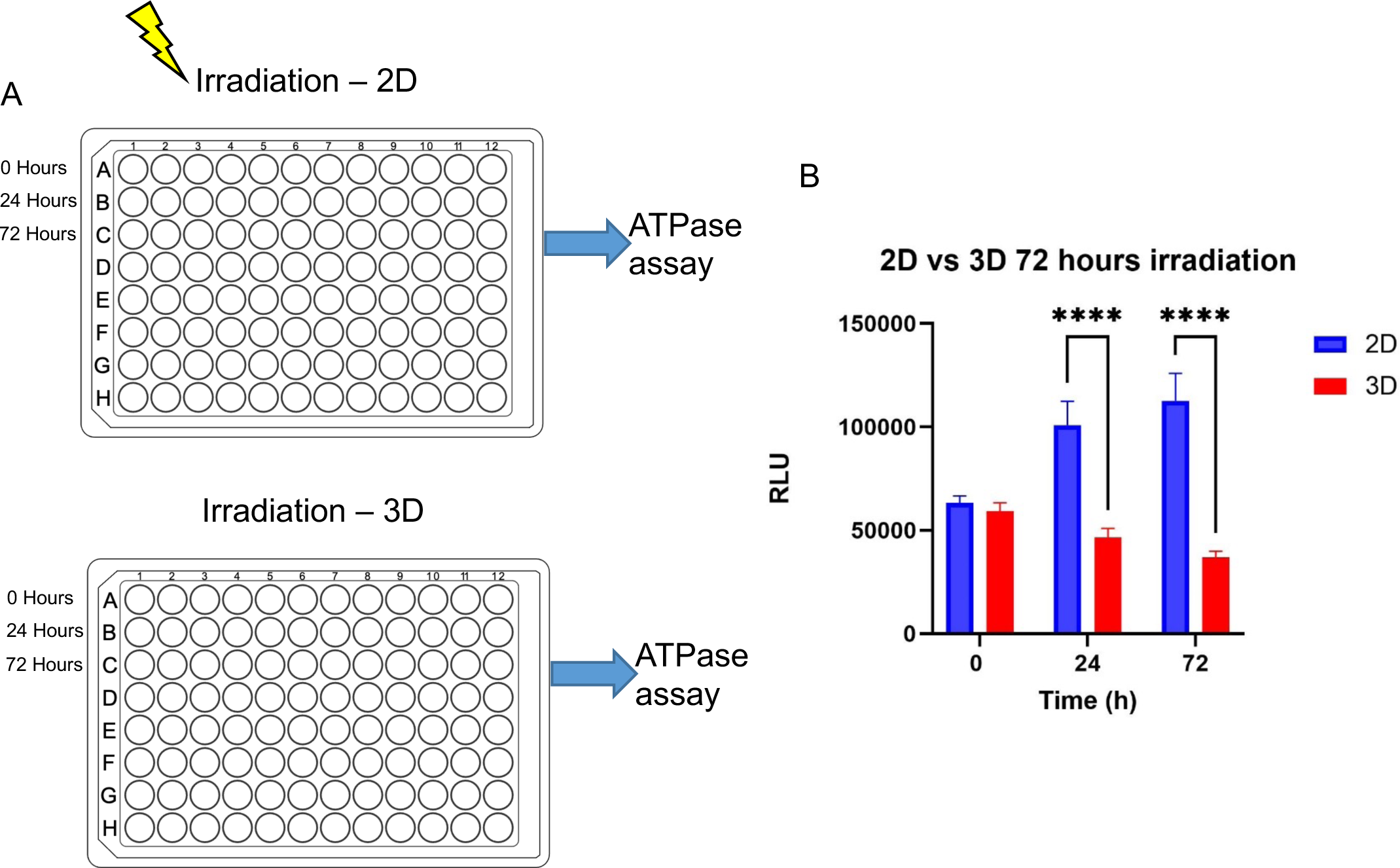
Response of HCC827 cells to irradiation differs depending on whether they are maintained in a 3D vs. 2D culture system. **(a**) Schematic of the irradiation experiment. ATPase luciferase assay was run to determine the viability of irradiated cancer cells at each time point. **(b)** Graphic representation of the ATP Cell Viability Luciferase Assay on live 2D vs. 3D cultures undergoing irradiation. Luminescence was captured with a luminometer and expressed as relative luminescence units (RLU). N=4 biological replicates. **** p<0.0001. Errors bars in SEM.

The results revealed significant differences in the response of the 2D and 3D cell cultures to irradiation. The 2D cell cultures continued to grow robustly, indicating their resistance to the treatment, whereas the 3D cell cultures exhibited decreased viability over a period of 72 hours, suggesting their increased sensitivity to irradiation (Figure 6B).

These findings provide evidence that the 3D cell culture configuration, compared to traditional monolayer cultures, may better reflect the response of lung cancer cells to a specific therapy, in this case, irradiation. This proof-of-concept study highlights the potential of 3D cell cultures as a more accurate model for predicting therapeutic efficacy in lung cancer patients, surpassing the limitations of existing tools.

### Comparison of 3D cultures to primary human lung cancer specimens

To assess the resemblance between the cell culture orientations (2D and 3D) and the primary EGFR adenocarcinoma tumor, we compared the bulk RNA seq data obtained from the 2D and 3D samples with the bulk RNA seq data of primary HCC827 lung tissue. Hierarchical clustering was performed on the differentially expressed genes. The heatmap demonstrated distinct clustering patterns, revealing a closer resemblance between the HCC827 primary tumor and the 3D cell culture system (Figure 7A). This suggests that the gene expression profiles of the 3D cell culture system more closely resemble those of the primary tumor compared to the 2D cell culture system.

**Figure 7:**
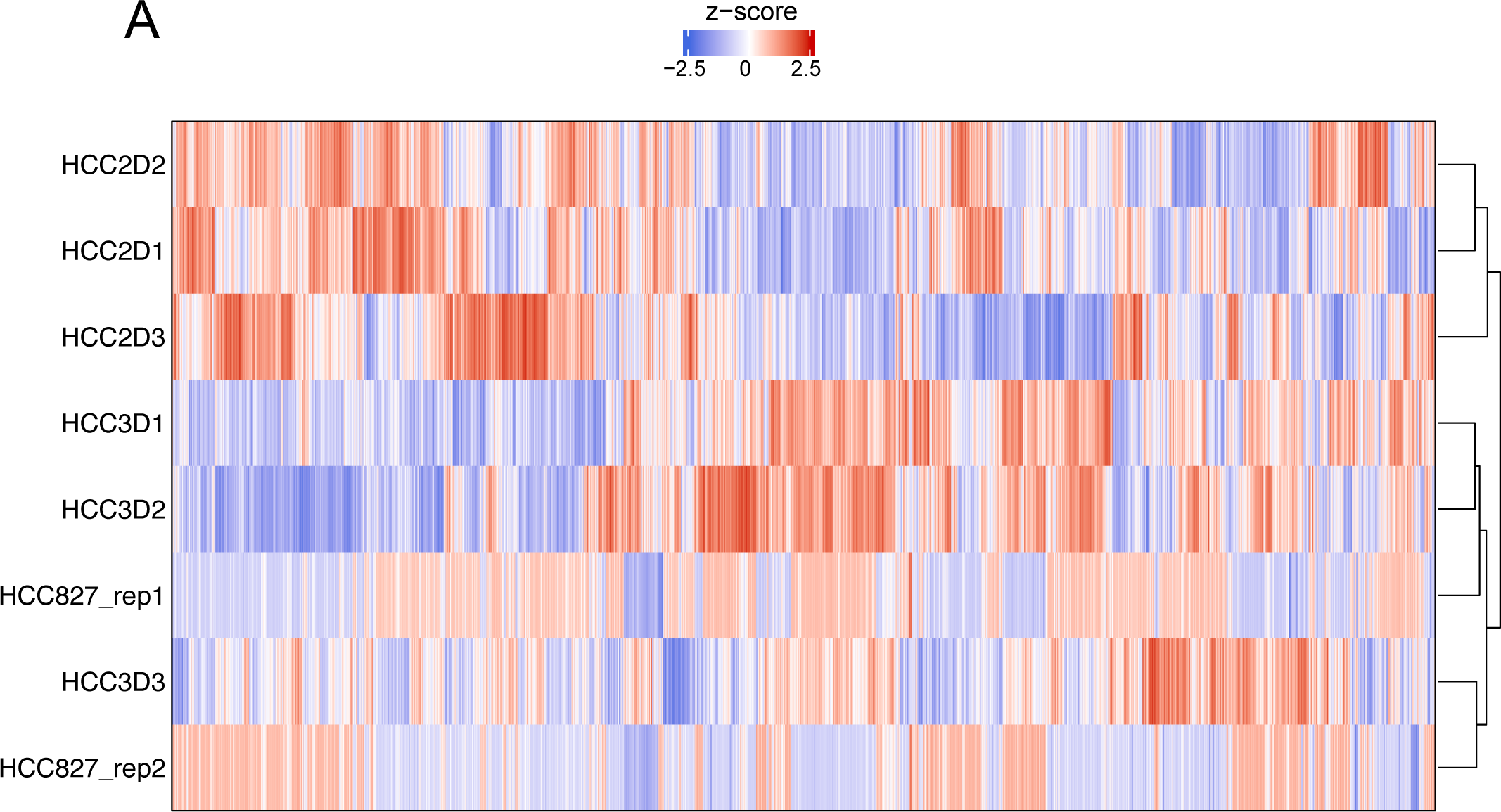
Comparison of 3D cultures to primary human lung cancer specimens Heat map analysis showing average expression (represented as z-score) of genes in the HCC827 2D cell cultures (HCC2Dx), 3D cell cultures (HCC3Dx) and primary human lung tumor (HCC827 repx). Hierarchical clustering is seen between the HCC827 primary tumor and the 3D cell culture system.

## DISCUSSION

This study highlights the clinical relevance of utilizing a three-dimensional (3D) model system for studying lung cancer development and therapeutics, compared to the traditional two-dimensional (2D) system. Our findings demonstrate that altering the cell culture configuration of a homogenous cancer cell line leads to significant differences in cell signaling, gene expression, and protein synthesis, which more closely resemble the characteristics of the actual tumor in situ. Notably, the 3D model system exhibited distinct gene expression patterns related to the tumor microenvironment, angiogenesis, and epithelial-mesenchymal transition. These gene expression changes were indicative of the complex interactions between cancer cells and their surrounding environment in the tumor microenvironment.

Furthermore, hierarchical gene clustering analysis confirmed that the 3D model system of EGFR adenocarcinoma closely resembled the primary human tumor, suggesting its superior representation compared to the 2D counterpart.

In addition to gene expression differences, we observed contrasting responses to irradiation and variations in the secretion of surfactant proteins, which plays a protective role in cell injury. These findings further emphasize the functional disparities between the 2D and 3D models, supporting the notion that the 3D model system better reflects *the in vivo* tumor environment.

The activation of oncogenic pathways was explored by analyzing gene expression related to the tumor microenvironment and the protein expression of phosphorylated ERK, a key component of the MAPK pathway. The 3D culture system of HCC827 cells exhibited upregulation of genes associated with tumor migration, invasion, metastasis, and proliferation, including the transcription factor *MYC*, which is known to play a crucial role in cell growth regulation. Additionally, the elevated protein expression of phosphorylated ERK in the 3D model system indicates enhanced activation of the MAPK pathway, highlighting the increased oncogenic signaling in the 3D model compared to the 2D model. Differential gene expression of cell growth and cell death-associated genes was also observed, with the 3D tumoroids displaying decreased expression of apoptotic molecule *CASP8* and necroptosis mediator *RIPK1*, while showcasing increased expression *MCM2*, a key regulator in DNA replication.

The 3D model system exhibited significant upregulation of genes related to cancer angiogenesis, indicating enhanced formation, maturation, and tone of blood vessels. Genes such as *VEGF* and *VEGFR*, known to induce the formation of new blood vessels, were particularly upregulated in the 3D system. This suggests that the epithelial cell-cell signaling in the 3D system promoted the development of new blood vessels. However, it was notable that HIF1A, typically secreted by cells in the hypoxic center of large tumors, showed decreased expression in the 3D model compared to the 2D system. This discrepancy might be explained by the fact that the 3D tumoroids did not reach a size large enough to generate a hypoxic center, while the 2D system became overgrown and produced a more hypoxic state. To determine if cell overgrowth influences HIF1A expression, experiments will need to be done at a lower confluency. Functional studies, although beyond the focus and scope of this paper, could be beneficial in further assessing the effects of angiogenesis activation. Ultimately, despite decreased HIF1A expression in the 3D tumoroids, the culmination of genes contributing to the activation of the angiogenesis pathway were significantly increased.

Furthermore, the EMT pathway genes were significantly upregulated in the 3D model system.

Vimentin and Fibronectin, molecules associated with a mesenchymal phenotype, showed higher expression in HCC827 cells cultured in 3D compared to 2D. Transitioning to a mesenchymal phenotype enhances the migratory ability of tumor cells. Additionally, there was a notable increase in the expression of matrix metallopeptidases, including *MMP9, MMP10, MMP11, MMP13, MMP14, MMP17*, and *MMP24*. These enzymes play a role in degrading the extracellular matrix, enabling cancer cells to migrate from their original site and potentially leading to invasion and metastasis. The invasive nature of the 3D tumoroids was evident as they expanded beyond the confines of the Matrigel and exhibited genotypic and phenotypic characteristics associated with increased mobility and invasiveness, indicating enhanced metastatic properties.

Overall, the 3D model system displayed enhanced angiogenesis, EMT pathway activation, and invasive properties compared to the 2D system. These findings underscore the relevance of utilizing the 3D model for studying cancer biology and therapeutic interventions, as it better captures the complex characteristics and behaviors of cancer cells in the context of their microenvironment.

The utilization of a 3D configuration of the same cells in our study also provided a better prediction of therapeutic responsiveness compared to the traditional 2D monolayers. Despite being the same cell type, the 2D monolayers continued to proliferate even after radiation exposure, whereas the 3D tumoroids exhibited a reduction in live cells. This observation, along with the increased hierarchical clustering of the 3D model with primary lung cancer tissue, suggests that the response of the 3D tumoroids to irradiation more closely aligns with the clinical response observed in patients.

By deliberately using the same adenocarcinoma cells in different configurations without the influence of non-cancer cells, we demonstrated that a 3D model that mimics the 3D orientation of cancer cells in their native environment better reflects their biology and treatment responsiveness. However, to further enhance the complexity and relevance of the 3D modeling, future studies should consider incorporating non-neoplastic cells that constitute the tumor microenvironment in situ, such as vascular cells, inflammatory cells, and extracellular matrix components. Additionally, it would be valuable to validate our conclusions using other types of lung cancers and additional adenocarcinoma cell lines to assess the generalizability of our findings.

In summary, our data strongly suggest that a 3D culture system of lung adenocarcinoma provides a more representative model of the primary tumor in terms of genotypic and phenotypic expression, as well as therapeutic responsiveness, compared to the traditional 2D culture system. The mere change in cell orientation from 2D to 3D resulted in differential gene expression of multiple cancer hallmarks, altered response to irradiation, and changes in surfactant protein expression.

These findings not only have implications for improving preclinical in vitro testing of therapeutics against organ system cancers but also offer insights into the origin and development of cancers within those organs when modeled as “tumoroids.”

## Acknowledgements

We thank Guillerma Garcia and the Sanford Burnham Prebys (SBP) Histology core for assisting in the staining procedures of the tumoroid samples and the SBP Bioinformatics Core for analysis. We thank Drs. Georgia Robins Sadler and Vanessa Malcarne for leading the program that made this research possible.

## Author Contributions

SLL and AE conceived, initiated, and coordinated the project. AE, SLL, RNM, and DF designed and performed the experimental work. SLL, DF, KV, EYS, and TM-C supplied reagents and assisted with data interpretation. AE, RM, DF, RNM, AB, BAG, ES, and SLL assisted in the development of figures. AE, RNM, EYS, and SLL wrote the manuscript and DF, RM, BAG, AB, ES, TM-C, HM, and KV edited. SLL, RM, SLL, RM, DF, TM-C, HM, RNM, KV, and EYS provided expertise and feedback.

