## Supplemental Figures 1-3 for "The effects of cell-cell orientation in modeling the hallmarks of lung cancer in vitro"

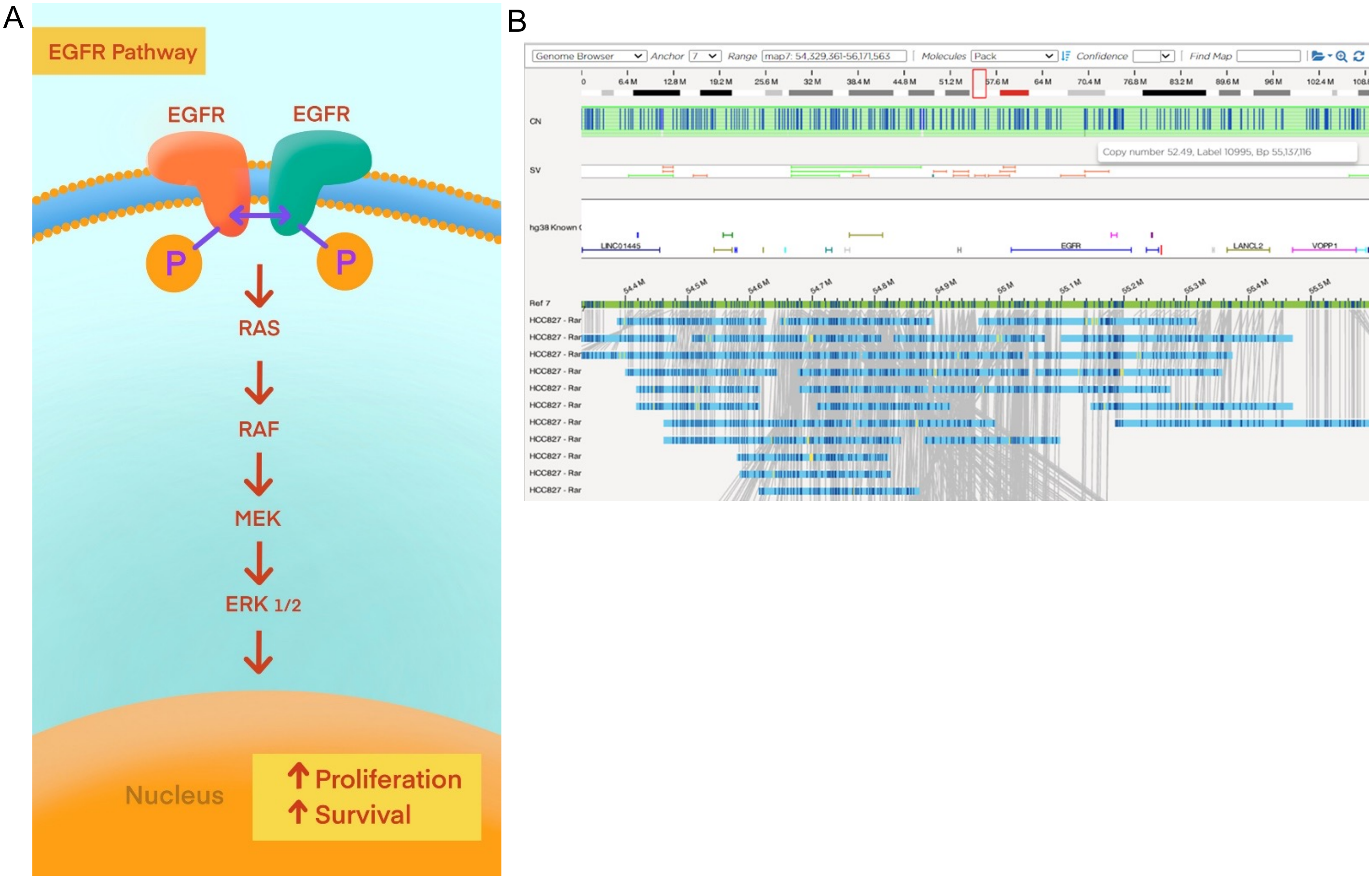


**Supplementary Figure 1: EGFR mutation in HCC827 cell line. (a)** Schematic showing the Intracellular RAS/RAF/MAPK signaling pathway after EGF receptor dimerization and activation in cancer cells. (b) Gene map showing the EGFR mutation of the adenocarcinoma cell line HCC827.

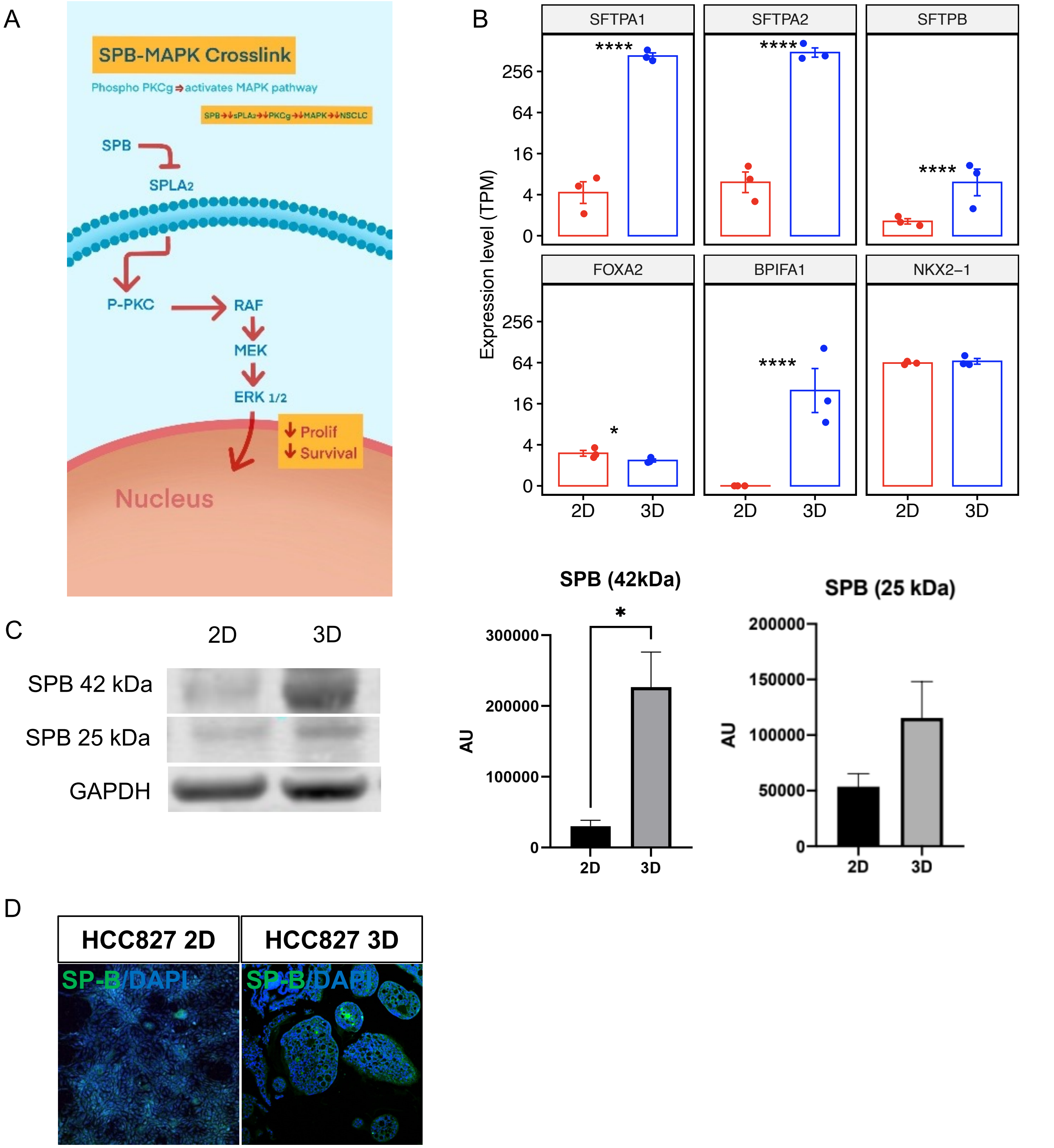


**Supplementary Figure 2: Surfactant protein B expression in 2D and 3D cell cultures**. (a) Schematic showing SPB interaction with the MAPK pathway. (b) TPM of surfactant-associated genes. (c) Western blot of SPB expression (42 kDa isoform and 25 kDa isoform) in HCC827 cells in 3D vs. 2D cultures (left panel). Quantification by densitometry (arbitrary units [AU]) of these cultures shows the that the same cells in 3D evince increased expression of SPB 42kDa isoform compared to the 2D cell cultures (right panel). N=3 biological replicates. *p<0.05. Error bars in SEM. (d) Immunostaining for SPB in 2D vs. 3D HCC827 cells. The data are representative of at least three independent experiments. A DAPI nuclear stain (blue) is used to visualize all cells in the field.

A

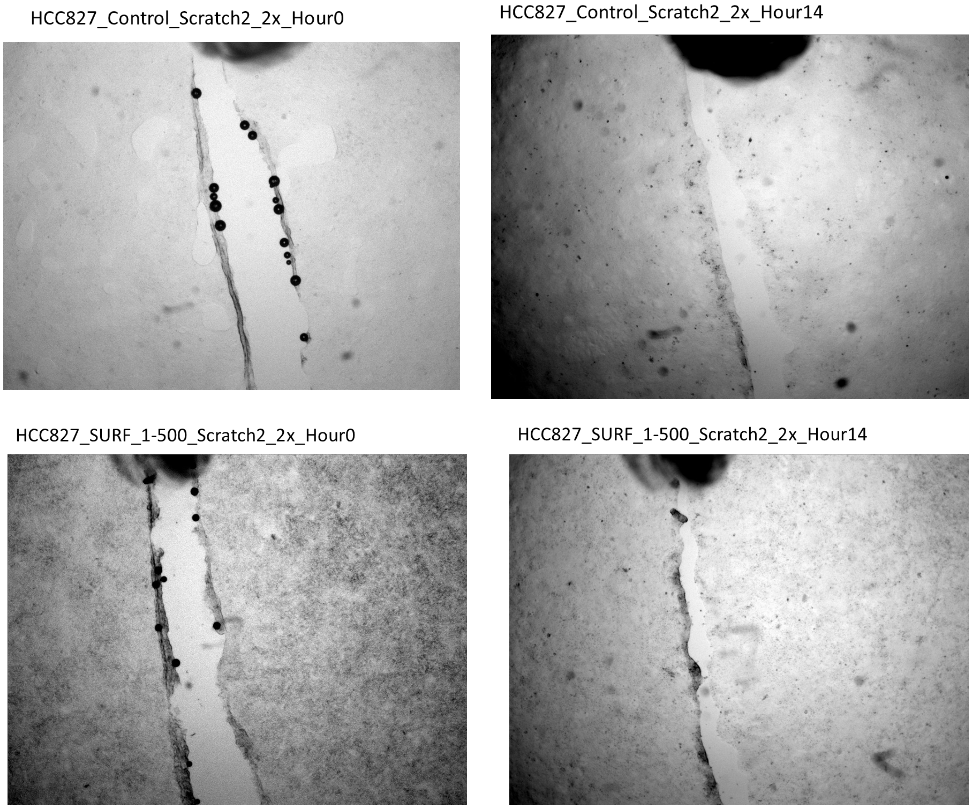


B

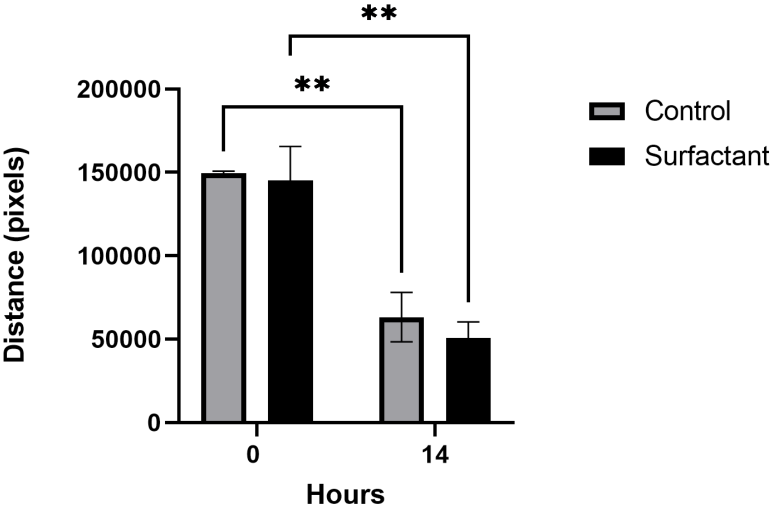


**Supplementary Figure 3: Effect of surfactant on wound healing in 2D cell cultures.** (a) Visual representation of the scratch performed in wells in the presence and absence of a whole lung surfactant emulsion. (b) Quantification of area healed over 14 hours.
